# A motor axonopathy in a mouse model of Duchenne Muscular Dystrophy

**DOI:** 10.1101/2020.01.22.914614

**Authors:** Justin S. Dhindsa, Angela L. McCall, Laura M. Strickland, Anna F. Fusco, Amanda F. Kahn, Mai K. Elmallah

## Abstract

Skeletal muscle weakness due to loss of dystrophin is a well-documented pathological hallmark of Duchenne muscular dystrophy (DMD). In contrast, the neuropathology of this disease remains understudied. Here, we characterize an axonopathy in the phrenic and hypoglossal (XII) nerves of *mdx* mice. We observe nerve dysfunction that we propose contributes to respiratory failure, the most common cause of death in DMD.

## Main

Duchenne muscular dystrophy (DMD) is a severe, progressive muscle wasting disorder that results from loss-of-function mutations in the *DMD* gene^1^. The encoded protein, dystrophin, is a large structural protein that connects the cytoskeleton of muscle fibers to the extracellular matrix via the sarcolemma, thereby acting as an important stabilizer of muscle during movement^2^. When dystrophin is deficient or absent, muscles are damaged during contraction, resulting in chronic inflammation and fibrosis^2,3^. Eventually muscle regeneration is inhibited, and healthy muscle cells are replaced by fibrotic and adipose tissue^3^.

In the most common mouse model of DMD – the *mdx* mouse – a point mutation in exon 23 of the dystrophin gene confers loss-of-function of the full-length dystrophin protein^4^. While the function of dystrophin and pathophysiology of DMD in skeletal muscle is well-described in this model, recent evidence suggests that dystrophin may also play an important in the nervous system^5^. Dystrophin deficiency in *mdx* mice alters neuron proliferation, survival, and/or differentiation as well as dysregulation of genes associated with axon development and synaptic organization^6,7^. In addition, there are significant morphological alterations have been observed in the neuromuscular junction (NMJ) of *mdx* mice, which hinder NMJ transmission and motor endplate function^8^.

Respiratory failure is the most common cause of death in patients with DMD^9^. Characterization studies of respiratory pathology in DMD have solely investigated muscular insufficiency and have overlooked potential neuropathologies^9–11^. In this study, we demonstrate that *mdx* mice exhibit a motor neuron axonopathy in the phrenic and hypoglossal (XII) nerves, which implies that an underlying neuropathology may also contribute to the characteristic respiratory failure of DMD. We show that *mdx* mice exhibit significant demyelination and loss of large-caliber axons. Furthermore, we find that mitochondria accumulate in the axoplasms of these nerves and exhibit morphological abnormalities. Altogether, we provide some of the first direct evidence that nervous system dysfunction may represent an important, but often overlooked, source of respiratory insufficiency in DMD.

Myelin ensheathment of axons is a vital process that allows for rapid impulse propagation along the nerve, promoting efficient nervous system function^12^. Myelination deficiency leads to axon degeneration and is a pathological hallmark of numerous neurological disorders^13^. Myelin sheath degradation – demyelination – is a well-documented phenotype in multiple sclerosis^14^, while myelin decompaction is a characteristic of ALS^15^. Therefore, we quantified the g-ratio (the ratio of the axon diameter to the diameter of the axon plus myelin) of axons with intact myelin in both nerves, as this ratio is a highly reliable indicator of optimal myelination^16^. Myelination is naturally optimized to achieve maximum efficiency in impulse propagation, and an abnormal g-ratio may suggest impaired nerve conduction^16^. For intact axons, *mdx* mice exhibit smaller g-ratio values in comparison to wildtype mice in both the phrenic and XII nerves, representing an axonopathy (**Fig. 1a**). Furthermore, axons that were not analyzed in *mdx* mice had severely collapsed and deformed myelin sheaths, exhibiting severe pathology (**Fig. 1f, g**). Together, these observations reflect a significant nerve pathology present in *mdx* mice.

**Fig. 1.**
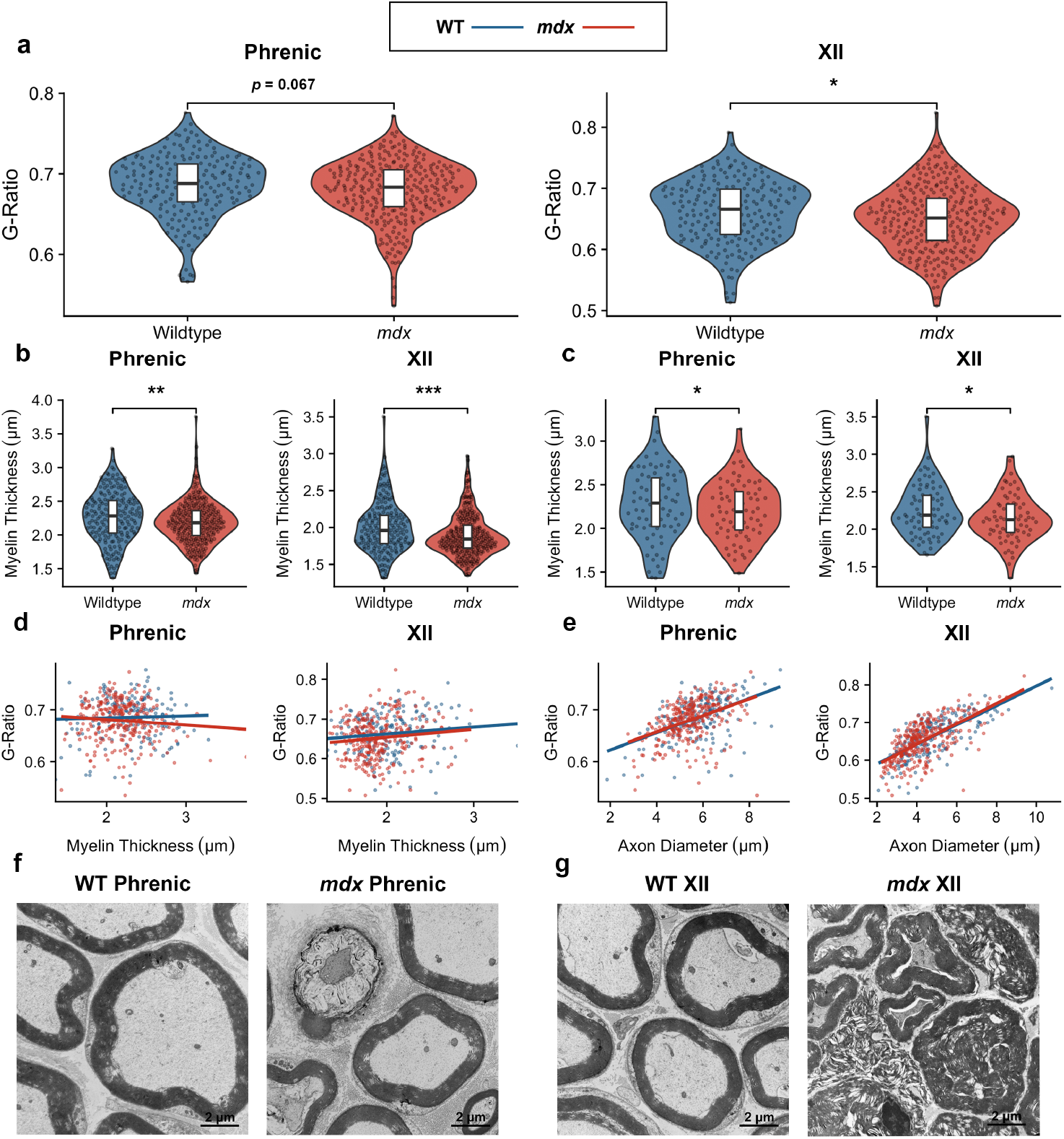
*mdx* mice experience suboptimal g-ratio values and demyelination. **a,** G-ratio values in phrenic and XII nerves from 12-month-old wild-type control mice (phrenic: 0.69 ± 0.003; XII: 0.66 ± 0.004; *n* = 200 fibers from two independent animals) compared with *mdx* mice (phrenic: 0.68 ± 0.002; XII: 0.65 ± 0.003; *n* = 300 fibers from three independent animals). **b,** Myelin sheath thickness in wild-type animals (phrenic: 2.27 μm ± 0.02; XII: 2.01 μm ± 0.02; *n* = 200 axons) compared with *mdx* mice (phrenic: 2.19 μm ± 0.03; XII: 1.89 μm ± 0.02; *n* = 300 axons). **c,** Myelin sheath thickness in only large-caliber axons (axon diameter > 5 μm) in wild-type mice (phrenic: 2.36 μm ± 0.02, *n* = 155 axons; XII: 2.25 μm ± 0.0, *n* = 76 axons) compared with *mdx* mice (phrenic: 2.28 μm ± 0.02, *n* = 195 axons; XII: 2.14 μm ± 0.04, *n* = 71 axons). **d,** Component of multivariate regression model, showing myelin sheath thickness against g-ratio in wild-type mice (phrenic: *p* = 1.48 × 10^−8^; XII: *p* = 9.12 × 10^−20^) and *mdx* mice (phrenic: *p* = 2.13 × 10^−13^; XII: *p* = 7.64 × 10^−18^). **e,** Component of multivariate regression model, showing axon diameter against g-ratio in wild-type mice (phrenic: *p* = 6.84 × 10^−21^; XII: *p* = 1.51 × 10^−48^) and *mdx* mice (phrenic: *p* = 6.23 × 10^−26^; XII: *p* = 2.32 × 10^−54^). **f, g,** Transmission electron microscopy images of intact myelin sheaths in wild-type mice, and collapsed and deformed myelin sheaths in *mdx* mice. Scale bars represent 2 μm.

Demyelination in *mdx* mice contributes to these observed g-ratio differences, as *mdx* mice exhibit significantly reduced myelin sheath thickness in both nerves (**Fig. 1b**). In addition, the size of the axons can impact the myelin thickness, as myelin thickness positively correlates with axon diameter. We find that larger-caliber axons (> 5 μm diameter)^17^ are selectively demyelinated in *mdx* mice, while smaller-caliber axons (< 5 μm diameter)^17^ are spared (**Fig. 1c, Fig. S1**). Because demyelination can lead to axon degeneration downstream^13^, we hypothesized that large-caliber axons are selectively degenerated in *mdx* mice. Furthermore, we considered that axon size may confound the g-ratio quantifications. To account for this potential confounder, we performed multivariate linear regression, regressing the g-ratio against both the axon diameter and myelin thickness (**Fig. 1d, e**). In both nerves, the models demonstrate that axon diameter is a better predictor of the g-ratio than myelin thickness. Therefore, we quantified the axon and fiber diameter (axon plus myelin diameter) and examined the distribution of axon caliber type.

In support of our hypothesis that large-caliber axons are selectively degraded, the axon and fiber diameters were noticeably reduced in *mdx* mice in both nerves (**Fig. 2a, b**). *mdx* mice clearly exhibited a skewed distribution of axons, favoring smaller axons and representing decreased heterogeneity of axon type (**Fig. 2c, d**). In contrast, wildtype mice show a more diverse composition of axon sizes, which is crucial for normal muscle function because large and small axons fulfill different roles. Small axons send high information signals to few muscle fibers, while large axons transmit low information signals to many muscle fibers^18^. Because large axons have a faster conduction time than small axons, they are vital for skeletal muscle function. Therefore, the selective loss of large-caliber axons in *mdx* phrenic and hypoglossal nerves likely hinders the proper function of the diaphragm and extrinsic tongue muscles, further exacerbating respiratory dysfunction in DMD.

**Fig. 2.**
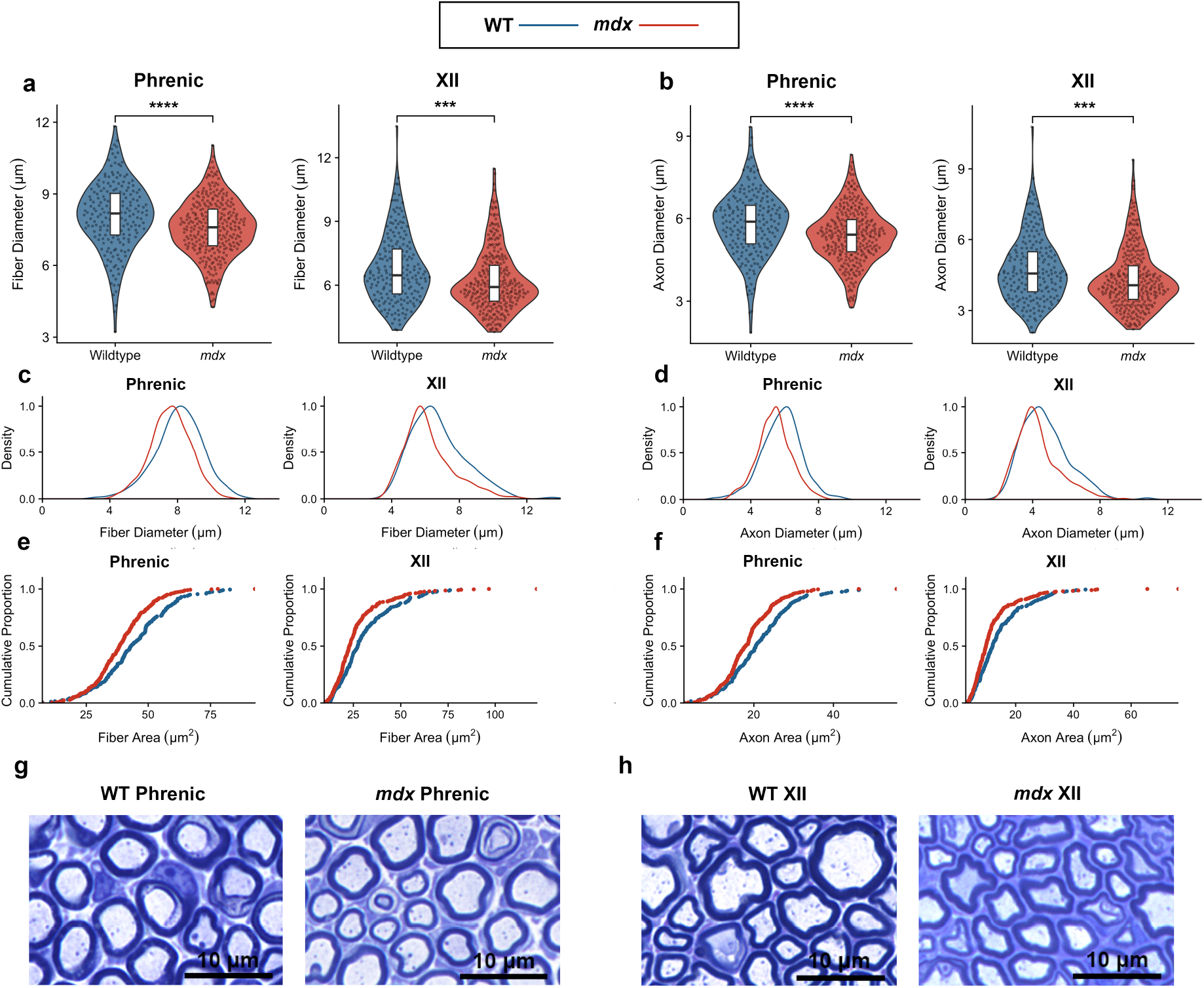
Large-caliber axons are selectively degraded in *mdx* mice. **a,** Fiber diameters (axon plus myelin diameter) in wild-type mice (phrenic: 8.08 μm ± 0.10; XII: 6.76 μm ± 0.11; *n* = 200 fibers) and *mdx* mice (phrenic: 7.58 μm ± 0.07; XII: 6.21 μm ± 0.08; *n* = 300 fibers). **b**, Axon diameter values in wild-type mice (phrenic: 5.81 μm ± 0.08; XII: 4.76 μm ± 0.10; *n* = 200 axons) and *mdx* mice (phrenic: 5.39 μm ± 0.06; XII: 4.32 μm ± 0.07; *n* = 300 axons). **c,** Distribution of fiber diameters in wild-type mice (phrenic SD: 1.41 μm; XII SD: 1.60 μm) and *mdx* mice (phrenic SD: 1.18 μm; XII SD: 1.44 μm). **d,** Distribution of axon diameters in wild-type mice (phrenic SD: 1.16 μm; XII SD: 1.37 μm) and *mdx* mice (phrenic SD: 0.10; XII SD: 1.26 μm). **e,** Cumulative proportion plots of fiber areas. **f,** Cumulative proportion plots of axon areas. **g, h,** Brightfield images of nerve cross sections. Scale bars represent 10 μm.

Mitochondrial dysfunction represents one of the earliest cellular deficits in *mdx* muscle^19^. Mitochondrial deficits impair muscle cells’ ability to repair damaged membranes, contributing to persistent myofiber damage^19^. Therefore, we employed transmission electron microscopy (TEM) to investigate whether mitochondrial abnormalities are also present in the axoplasms of phrenic and XII nerves. The axoplasm – the cytoplasm of the axon – serves as an avenue for the transport of proteins, mRNA, lipids, and mitochondria in both the anterograde and retrograde directions^20^. We observe that the number of mitochondria in the axoplasmic space is significantly increased in both nerves of *mdx* mice, and that there is a significant increase in mitochondrial diameter in only the XII nerve of *mdx* mice (**Fig. 3b, c)**. Additionally, the density of mitochondria is increased in both nerves of *mdx* mice (**Fig. 3d)**. The increase in mitochondrial diameter indicates swelling, which is a hallmark of dysfunction of these organelles^21^. Furthermore, the increased number and density of mitochondria suggests accumulation, which can represent either reduced axonal transport or impaired mitophagy^22^. Both of these features have been associated with other neuromuscular diseases, such as Charcot-Marie-Tooth type 2A^22^.

**Fig. 3.**
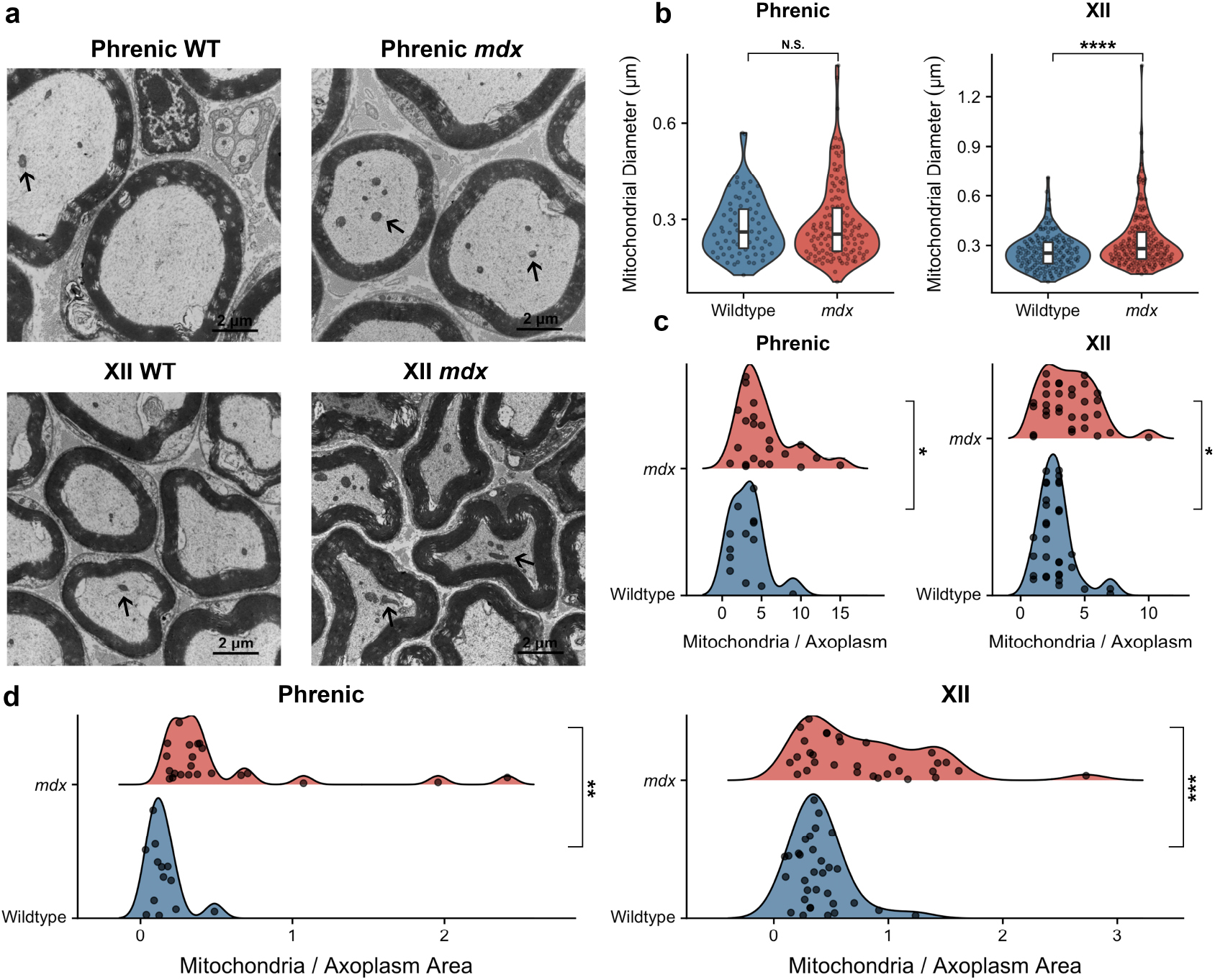
*mdx* mice exhibit altered morphology and accumulation of mitochondria. **a,** TEM cross-sections of wild-type and *mdx* mouse nerves. Black arrows pointing to mitochondria. Scale bars represent 2 μm. **b,** Mitochondrial diameter in wild-type mice (phrenic: 0.28 μm ± 0.01, *n* = 70 mitochondria; XII: 0.26 μm ± 0.01, *n* = 145 mitochondria) and *mdx* mice (phrenic: 0.29 μm ± 0.01, *n* = 127 mitochondria; XII: 0.33 μm ± 0.01, *n* = 173 mitochondria). **c,** number of mitochondria per axoplasm in wild-type mice (phrenic: 3.23 ± 0.62, *n* = 13 axoplasms; XII: 2.78 ± 0.25, *n* = 32 axoplasms) and *mdx* mice (phrenic: 5.41, ± 0.76, *n* = 22 axoplasms; XII: 3.77 ± 0.37, *n* = 31 axoplasms). **d,** Density of mitochondria in axoplasms in wild-type mice (phrenic: 0.15 ± 0.03; XII: 0.40 ± 0.04) and *mdx* mice (phrenic: 0.54 ± 0.12; XII: 0.81 ± 0.10).

Proper myelination is required for maintenance of normal axon transport and long-term survival^23^, so the axonal demyelination we observe may have consequences on axonal transport leading to mitochondrial accumulation. Alternatively, axons may be transporting more mitochondria to the NMJ as a compensatory response to NMJ degradation in *mdx* mice^8^. Because axon transport deficits are linked with motor neuron death and dysfunction^20^, we quantified the number of phrenic and XII motor neurons (**Fig. S2**). There was no significant difference between the number of neurons between wildtype and *mdx* mice for both phrenic and XII neurons, illustrating that motor neurons are not degraded in *mdx* mice. Therefore, the pathology at this time point seems to be constrained to the axons and does not affect the soma.

Taken together, our data suggest significant neuropathologies in the phrenic and XII nerves of *mdx* mice. However, because full-length dystrophin is not expressed in the spinal cord of healthy mice or humans^24^, we propose that these pathologies are secondary to the characteristic muscle-wasting. Specifically, we speculate that the nerve dysfunction may be a consequence of dystrophin deficiency at the NMJ. Dystrophin accumulates at the postsynaptic membrane in the form of the dystrophin-associated glycoprotein complex (DGC), and deficiency of the DGC has been shown to disrupt both the pre- and postsynaptic structures of the NMJ in the *mdx* model^25^. Therefore, we suspect that pathology starts at the NMJ and then acts in a retrograde manner later affecting the nerves, a process termed “dying-back”^26^.

Overall, our data strongly suggest that there are striking neuropathologies involved in DMD that often go overlooked. We find that the phrenic and XII nerves display key pathological features that are characteristic of other neuromuscular diseases but have previously gone uncharacterized in DMD. These findings contribute to the emerging evidence that dystrophin deficiency, induced by *DMD* mutations, adversely affects the nervous system in addition to muscle fibers. Further investigation of potential neuropathologies must be done to fully elucidate the disease mechanisms of DMD. Additionally, our study has important implications for future treatment of DMD because correcting the muscle pathology alone will likely not completely ameliorate the respiratory deficiency associated with DMD.

## Acknowledgements

The authors thank the members of the Duke University Electron Microscopy Core, Sara Miller, PhD, Ricardo Vancini, PhD, and Harold Makeel, for their expertise and preparation of tissues.

## Author Contributions

Conceptualization and Experimental Design: JSD, MKE. Experimental Execution and Analysis: JSD, ALM, LMS, AFF, AFK. Manuscript Writing: JSD. Editing: JSD, MKE. Supervision: MKE.

## Methods

### Experimental Animals

All mice were approved by the Duke University Institutional Animal Care and Use Committee (IACUC) under protocol A233-171-10. C57Bl6/J, wildtype (WT), and C57BL/10ScSn-Dmd^*mdx*^/J (*mdx*), mice were obtained from the Jackson Laboratory and housed at the Duke University Division Laboratory Animal Resources.

### Nerve Processing and Images

Phrenic and hypoglossal (XII) nerves were harvested from 12-month-old wildtype (*n* = 2) and *mdx* mice (*n* = 3). The nerves were placed in 2.5% glutaraldehyde and 0.1% sodium cacodylate. They were then processed, embedded in hard plastic, sectioned to 1 μm and consequently stained with 1% toluidine blue and 1% sodium borate by the Duke University Electron Microscopy Core. Semi-thin sections were imaged on a Keyence BZ-X710 All-in-One Fluorescence Microscope. Light micrographs were analyzed using a public downloadable ImageJ plugin^27^ to examine the g-ratio, fiber diameter, axon diameter, and myelin thickness. The g-ratio was determined to be the diameter of the inner axon to the total outer diameter of the axon plus the myelin sheath^16^. 100 randomly selected axons from each nerve were manually outlined for each animal.

For electron microscopy, phrenic and hypoglossal (XII) nerves from wildtype (*n* = 2) and *mdx* (*n* = 2) mice were placed in 2.5% glutaraldehyde and 0.1% sodium cacodylate and then post-fixed in 1% osmium tetroxide. The nerves were placed in 1% uranyl acetate and then dehydrated with acetone. They were then processed in epoxy resin (EPON), cut into 60 nm ultrathin sections on a Reichert Ultracut E ultramicrotome, and stained with 2% uranyl acetate and SATO’s Lead stain. The nerves were imaged on a Philips CM12 electron microscope. The area of the axoplasm and diameter of mitochondria was measured using ImageJ 1.48v.

### Motor Neuron Counting

12-month-old mice were anesthetized and harvested for their spinal cords, which were fixed in 4% paraformaldehyde in phosphate-buffered saline (PBS). The spinal cords were placed in 30% sucrose at 4°C. The spinal cord was separated into sections: medulla, cervical spinal cord, thoracic and lumbar. The sections were embedded in OCT and frozen at −80°C before cryosectioning. Medulla and cervical spinal cords were cut into 40 μm cross sections using a Leica CM3050 S Cryostat. The sections were preserved in a 96-well plate in 2% PFA. Every third cervical spinal cord cross-section, and every other section of the medulla were transferred to a master 96-well plate with 1x PBS for histochemical analysis.

The mounted tissues were then stained using Cresyl Violet. The Cresyl Violet solution was prepared using 0.1% Cresyl Violet Acetate in distilled H_2_O and stirred overnight The slides were sequentially submerged in the following solutions: distilled H_2_O, Cresyl Violet, distilled H_2_O, 50% EtOH, 75% EtOH, 95% EtOH, 100% EtOH, Xylene, and then re-submerged in Xylene. Sections were then cover slipped and imaged using a Leica DMRA2 for brightfield microscopy. The images were analyzed and counted by two different individuals, and their results were compared with each other.

### Statistical Analysis

Differences between wild-type and *mdx* mice were evaluated using an unpaired *t*-test. Significance was set at *p* < 0.05 (*); *p* < 0.01 (**); *p* < 0.001 (***); *p* < 0.0001 (****). Values in figure legends are reported as the mean ± standard error of mean (SEM). Computations were completed using RStudio, an integrated development editor of R. Plots were also generated in RStudio.

