## Supplemental Figures 1 and 2 for "A motor axonopathy in a mouse model of Duchenne Muscular Dystrophy"

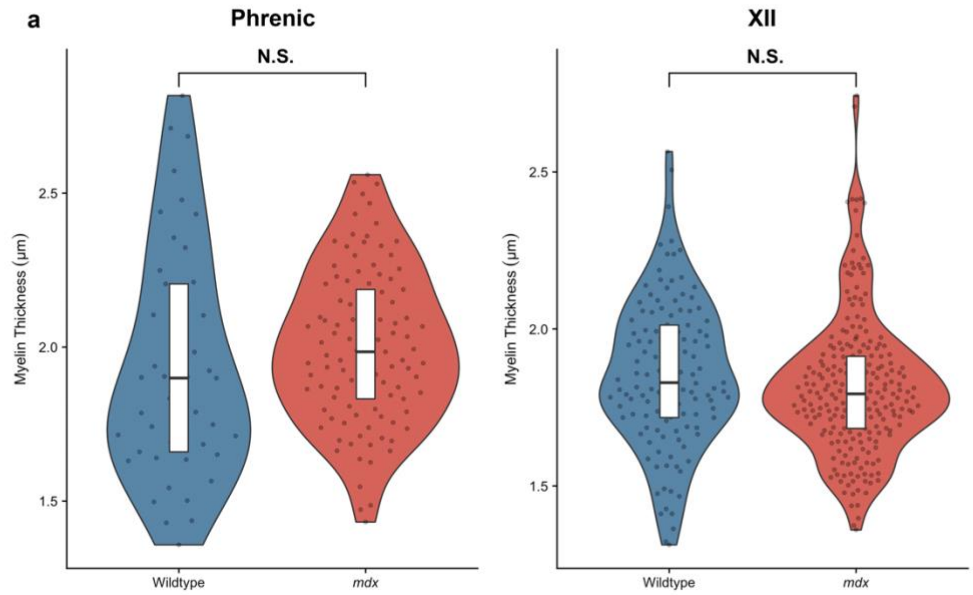


**Fig. S1 Small-caliber axons do not experience demyelination in *mdx* mice**
**a,** Myelin sheath thickness of small-caliber axons (< 5 μm) in wild-type mice (phrenic: 1.94 μm ± 0.06, *n* = 45 axons; XII: 1.86 μm ± 0.02, *n* = 124 axons) and *mdx* mice (phrenic: 2.00 μm ± 0.02, *n* = 105 axons; XII: 1.82 μm ± 0.01, *n* = 229 axons)

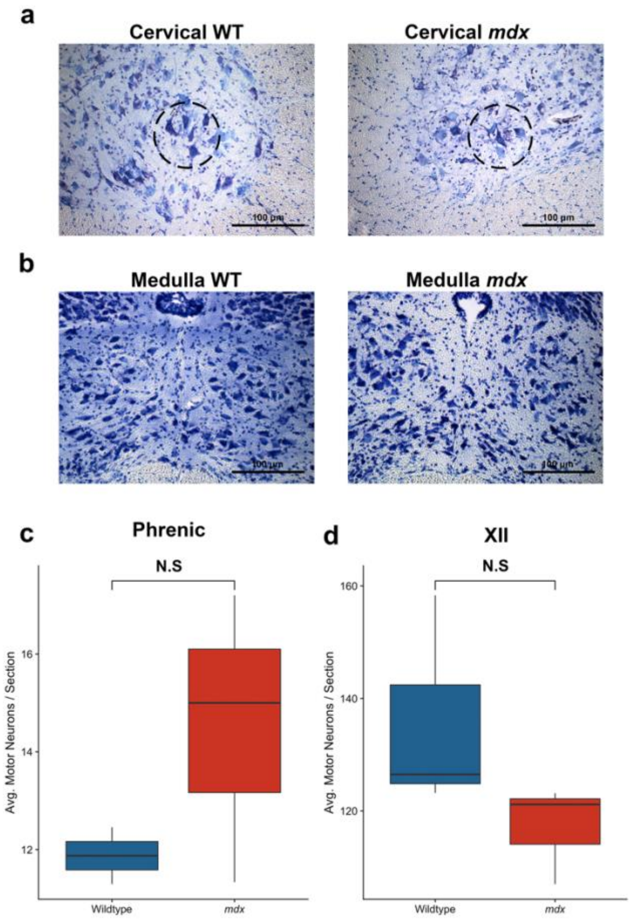


**Fig. S2 Phrenic and XII motor neuron counts**

**a,** Bright field images of C4 spinal cord sections. Dashed circle around the phrenic motor pool. **b,** Bright field images of medulla sections. Scale bars represent 100 μm. **c,** Average phrenic motor neurons per cervical spinal cord section (Wildtype: 11.87 motor neurons ± 0.58, *n* = 2 mice; *mdx:* 14.51 motor neurons ± 1.71, *n* = 3 mice). **d,** Average XII motor neurons per medulla section (Wildtype: 136.01 motor neurons ± 11.20, *n* = 3 mice ; *mdx:* 117.10 motor neurons ± 5.08, *n* = 3 mice).
